# Dominant nitrogen metabolisms of a warm, seasonally anoxic freshwater ecosystem revealed using genome resolved metatranscriptomics

**DOI:** 10.1101/2023.08.22.554355

**Authors:** J. M. Fadum, M. A. Borton, R. A. Daly, K. C. Wrighton, E. K. Hall

**Affiliations:** Graduate Degree Program in Ecology, Colorado State University, Fort Collins, CO, USA; Department of Ecosystem Science and Sustainability, Colorado State University, Fort Collins, CO, USA; Department of Soil & Crop Sciences, Colorado State University, Fort Collins, CO, USA

## Abstract

Reactive nitrogen (N) is one of the principal drivers of primary productivity across aquatic ecosystems. However, the microbial communities and emergent metabolisms which govern N cycling in tropical lakes are both distinct from and poorly understood relative to those found in temperate lakes. This latitudinal difference is largely due to the warm (>20 °C) temperatures of tropical lake anoxic hypolimnions (deepest portion of a stratified water column) which result in unique anaerobic metabolisms operating without the temperature constraints found in lakes at temperate latitudes. As such, tropical hypolimnions provide a platform for exploring microbial membership and functional diversity. To better understand N metabolism in warm anoxic waters, we combined measurements of geochemistry and water column thermophysical structure with genome resolved metatranscriptomic analyses of the water column microbiome in Lake Yojoa, Honduras. We sampled above and below the oxycline in June 2021, when the water column was stratified, and again at the same depths and locations in January 2022, when the water column was mixed. We identified 335 different lineages and significantly different microbiome membership between seasons and, when stratified, between depths. Notably, *nrfA* (indicative of dissimilatory nitrate reduction to ammonium) was upregulated relative to other N metabolism genes in the June hypolimnion. This work highlights the taxonomic and functional diversity of microbial communities in warm and anoxic inland waters, providing insight into the contemporary microbial ecology of tropical ecosystems as well as inland waters at higher latitudes as water columns continue to warm in the face of global change.

**Importance:** In aquatic ecosystems where primary productivity is limited by nitrogen (N), whether continuously, seasonally, or in concert with additional nutrient limitations, increased inorganic N availability can reshape ecosystem structure and function, potentially resulting in eutrophication and even harmful algal blooms (HABs). Whereas microbial metabolic processes such as mineralization and dissimilatory nitrate reduction to ammonium (DNRA) increase inorganic N availability, denitrification removes bioavailable N from the ecosystem. Therefore, understanding these key microbial mechanisms is critical to the sustainable management and environmental stewardship of inland freshwater resources. This study identifies and characterizes these crucial metabolisms in a warm, seasonally anoxic ecosystem. Results are contextualized by an ecological understanding of the study system derived from a multi-year continuous monitoring effort. This unique dataset is the first of its kind in this largely understudied ecosystem (tropical lakes) and also provides insight into microbiome function, and associated taxa, in warm anoxic freshwaters.

## Introduction

Aquatic biogeochemistry is defined by the microbial consortia that inhabit these diverse and rapidly changing ecosystems. However, our understanding of microbially driven biogeochemical transformations and associated taxonomies in inland waters is heavily skewed towards temperate ecosystems. Low latitude aquatic microbiomes are comparatively poorly characterized and understood, despite the demonstrable importance of microbial food webs in tropical lakes (1) and the contributions of diverse microbial metabolisms to local and global biogeochemical cycles (2-5). This geographic disparity presents a significant barrier to understanding the ways in which microbial communities shape ecosystem function and is highlighted by the frequency of unclassified OTUs reported in assessments of tropical lake microbial community composition (6).

Seasonal anoxia occurs across morphologically diverse lake ecosystems, including those at both temperate and tropical latitudes. However, biogeochemical cycling in tropical lakes which maintain a stratified water column (either permanently or seasonally) may be particularly distinct from that in their temperate counterparts, in part, due to warm (>20 °C) temperatures of their anoxic waters (compared to ∼4 °C at temperate latitudes) (7). Understanding emergent metabolisms when microbial communities are released from temperature controls which constrain anaerobic metabolisms at high latitudes offers critical insight into the microbial ecology and biogeochemistry of contemporary ecosystems as well as potential functional shifts under future climate scenarios. Though recent work has gained insight into the microbial ecology of permanent anoxic zones in the open ocean (8-11), and in anoxic waters of temperate and arctic lakes (12-14), microbial mediated biogeochemistry of warm inland waters that sustain anoxia for all, or parts, of the year remain less well described, with some notable exceptions (15, 16).

Understanding the microbial drivers of nitrogen (N) biogeochemistry in aquatic ecosystems is particularly important because reactive N is one of the principal drivers of primary productivity in inland waters. One important source of reactive N to surface waters in seasonally stratified tropical lakes is the hypolimnion (the deepest layer of a stratified water column) due to accumulation of reactive N during stratification and release of that reactive N to the photic zone during turnover (17-21). However, the anaerobic metabolisms (and associated taxa) which contribute to this reactive N accumulation are poorly defined. Therefore, to empirically identify which microbial pathways drive N biogeochemistry in warm anoxic water columns and provide a mechanistic explanation of reactive N accumulation in tropical hypolimnions, we provide the first metagenome resolved metatranscriptomic analysis of a tropical lake water column under both oxic and anoxic conditions. We hypothesized that, in addition to mineralization of organic N, dissimilatory nitrate reduction to ammonium, (DNRA) may be a unique feature of warm anoxic hypolimnions and an important contributor to the observed accumulation of NH_4_^+^ in the anoxic strata of the water column of our study site, Lake Yojoa.

Lake Yojoa (∼ 83 km^2^ surface area, 1.4 km^3^ volume, 27.3 m annual average max depth) is located in the center of a ∼337 km^2^ mixed land use/landcover watershed in West-Central Honduras. The lake supports natural fisheries in addition to one large industrial aquaculture operation. The watershed has persistently warm temperatures (annual average air temperatures above 20 °C) and receives approximately 2 m of annual precipitation with the warmest months typically corresponding with the monsoon season (June to October). Primary productivity during the mixed water column phase is largely driven by hypolimnetic nutrients which are released to the photic zone following water column mixing, typically in November (17, 22).

## Results

### Water column redox state and nutrient chemistry

To establish seasonal differences in redox conditions throughout the water column, we compared DO concentrations in surface and deep waters in June 2021 and again in January 2022. In June, Lake Yojoa was stratified with an oxycline at ∼12 m depth (which only varied slightly among the three sampling stations, Fig. S1) (Table 1.) Therefore, the samples collected in June at 16 m were the only samples taken from an anoxic portion of the water column while all other samples (June surface and January at both depths) came from oxic waters. Although the June hypolimnion was the only reduced environment sampled, temperature was the same as in January across all depths (Table 1). Temperature above 12 m in June significantly (*p*< 0.001) exceeded temperature in June below the oxycline and at both depths in January.

**Table 1.** Thermophysical structure of Lake Yojoa during sampling events (mean of three sampling locations ± standard error).

|  | June 2021<br>DO (mgL<sup>-1</sup>)<br>(n=) | June 2021<br>Temperature (°C)<br>(n=) | January 2022<br>DO (mgL<sup>-1</sup>)<br>(n=) | January 2022<br>Temperature (°C)<br>(n=) |
| --- | --- | --- | --- | --- |
| Depth < 12 m | 6.82 ± 0.23<br>(n=18) | 28.32 ± 0.14<br>(n=18) | 5.51 ± 0.19<br>(n=18) | 24.40 ± 0.02<br>(n=18) |
| Depth > 12 m | 0.00 ± 0.00<br>(n=12) | 24.56 ± 0.17 (n=12) | 5.14 ± 0.22<br>(n=12) | 24.32 ± 0.02<br>(n=12) |

To assess seasonal and spatial differences in nutrient chemistry, we compared the two dominant forms of reactive N (NH_4_^+^ and NO_3_^-^), TP, and DOC between depths and seasons. In January, when the water column was mixed, there was little variation in measured geochemical parameters between the two sampling depths (Fig. 2A-D, Table S1). In contrast, in June, when the water column was stratified, we observed significant (*p*<0.05) differences in multiple geochemical parameters between the surface and hypolimnion (Fig. 2 E-H, Table S1). In particular, hypolimnetic concentrations of NH_4_^+^ were much greater than surface concentrations of NH_4_^+^ at all three stations (*p*= 0.001), though differences varied across stations (Fig. 2E). TP was relatively low (< 3 µM) across all locations and depths in June and differences between surface and hypolimnetic concentrations were not significant (Fig. 2F). Similarly, NO_3_^-^ was also low in both the hypolimnion and the surface, and not significantly different between strata for all three stations (Fig. 2H). DOC concentrations, though marginally higher in the surface than in the hypolimnion, did not have as pronounced differences as did NH_4_^+^(*p*= 0.04) (Fig. 2G).

Geochemical differences in June were primarily characterized by pronounced differences in DO and NH_4_^+^ and to a lesser extent TP between strata, whereas geochemistry in January was largely homogenous between the two depths.

We also identified less pronounced but still significant differences in geochemistry within depths among our three sampling stations (Fig. 1). In January, there were few significant differences among sampling locations at the same depth with the exception of NH_4_^+^, which was significantly (*p*= 0.0008 at 1 m and *p*= 0.0003 at 16 m) greater at the northern most sampling location at both depths (48.00 ± 0.40 µM at 1 m and 50.20 ± 0.10 µM at 16 m) compared to the central (34.00 ± 1.01 µM at 1 m and 37.05 ± 0.95 µM at 16 m) and southern location (32.05 ± 0.05 µM at 1 m and 32.35 ± 0.05 µM at 16 m) (Fig. 2A). In June, hypolimnetic NH_4_^+^ was significantly higher at the central station compared to the northern and southernmost stations (*p*= 0.02, Tukey’s HSD, 95% confidence interval). Hypolimnetic TP was also significantly higher at the northernmost station (in comparison to the southern station, *p* = 0.009, and in comparison to the central station, p=0.014). Surface TP was only significantly different at the southernmost station (*p* = 0.03). Therefore, while there were quantifiable spatial differences in nutrient chemistry, we did not identify consistent patterns that suggested any particular station was consistently enriched in both reactive N and TP compared to other sampling stations.

**Figure 1.**
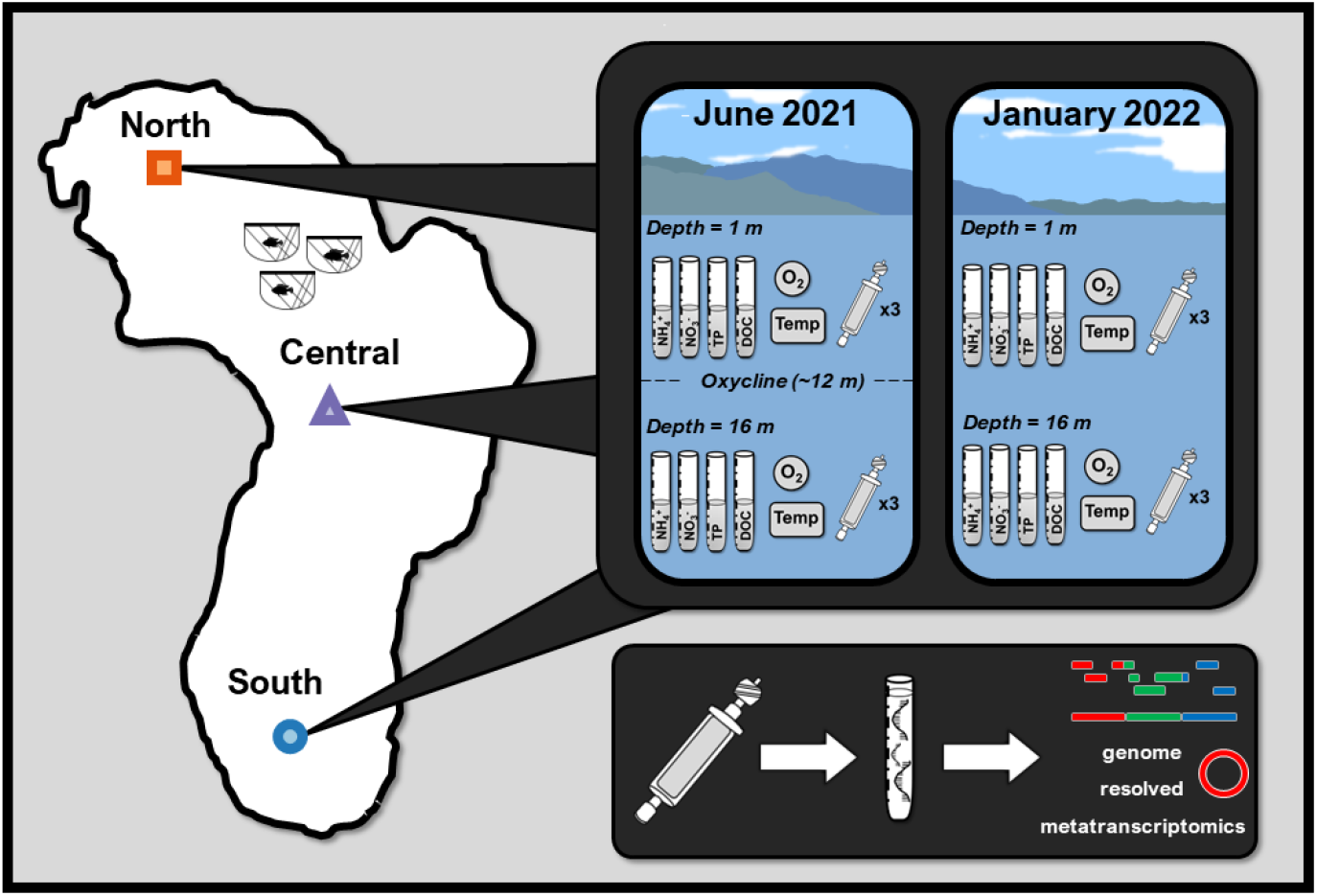
Data collection conceptual figure identifying the three pelagic sampling locations (North, Central and South) and the types of samples collected (geochemical, thermophysical structure, and metatranscriptome-in triplicate) followed by extraction, sequencing, genome assembly and metatranscriptomic mapping to assembled genome.

**Figure 2.**
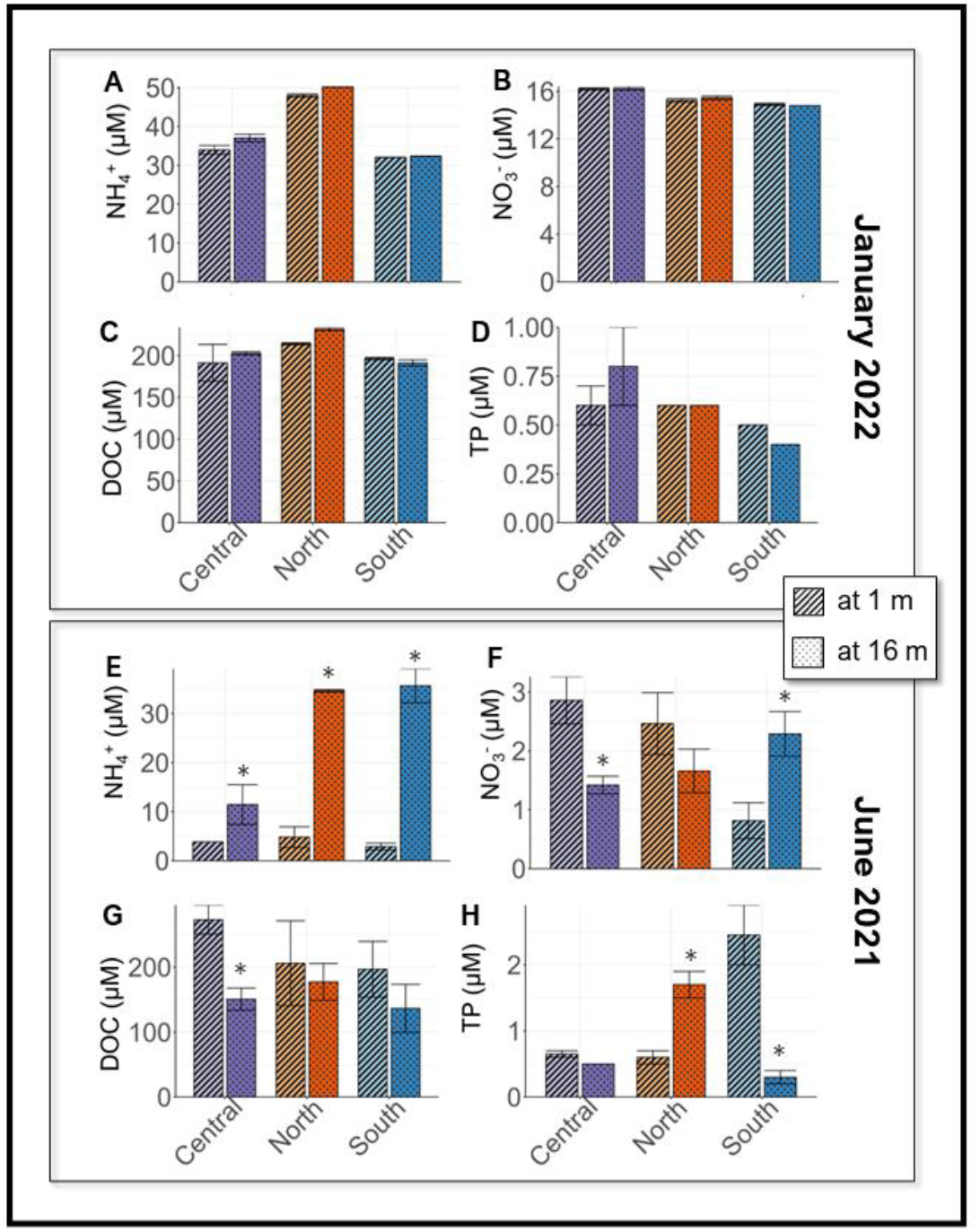
Nutrient concentrations in Lake Yojoa at 1 and 16 m across three sampling locations, mean ± SE, in January 2022 **(A-D)** and June 2021 **(E-H)**. Significant differences between 1 m and 16 m identified with an asterisk (*).

### Microbiome membership in time and space

Our dereplicated database was composed of 572 MAGs. However, in order to focus on active biogeochemical pathways, we limited our assessment of microbiome membership to the MAGs detected in the metatranscriptomic data (n=552). Within the Lake Yojoa metatranscriptome, we identified 335 different lineages (76 classes across 37 phyla). Many genomes represent taxonomies undefined at the family (4.5%), genus (21.1%), or species (86.5 %) level. Lineages with novel families often followed unnamed orders (n= 3) or alphanumerically identified orders (n=13). The abundance of novel lineages in our dataset highlights the under-representation of tropical freshwater ecosystems in public genome databases and the degree to which tropical lake microbiomes are still poorly described (6).

The phyla *Proteobacteria* (n= 97), *Bacteroidota* (n=90), *Planctomycetota* (n=70), *Verrucomicrobiota* (n=66), and *Actinobacteriota* (n= 56) dominated the represented lineages, as may be expected for freshwater lake ecosystems (23) (Fig. 3A). In June 2021, surface waters were dominated with *Lyngbya robusta* and *Microcystis wesenbergii*, two common cyanobacteria (24, 25) as expected under conditions of N limitation (17). C*yanobacteriia* was the most abundant class during both seasons and across both depths. Interestingly, phyla more frequently identified in marine or saline environments, such as *Thermoproteota* and *Halobacteriota* were also identified. Additionally, taxonomies associated with gut microbiomes including *Fibrobacterota, Fusobacteriota*, and *Firmicutes*, were identified, suggesting organic pollution from municipalities, ranching, and/or the industrial aquaculture operation located within the north basin of the lake.

**Figure 3.**
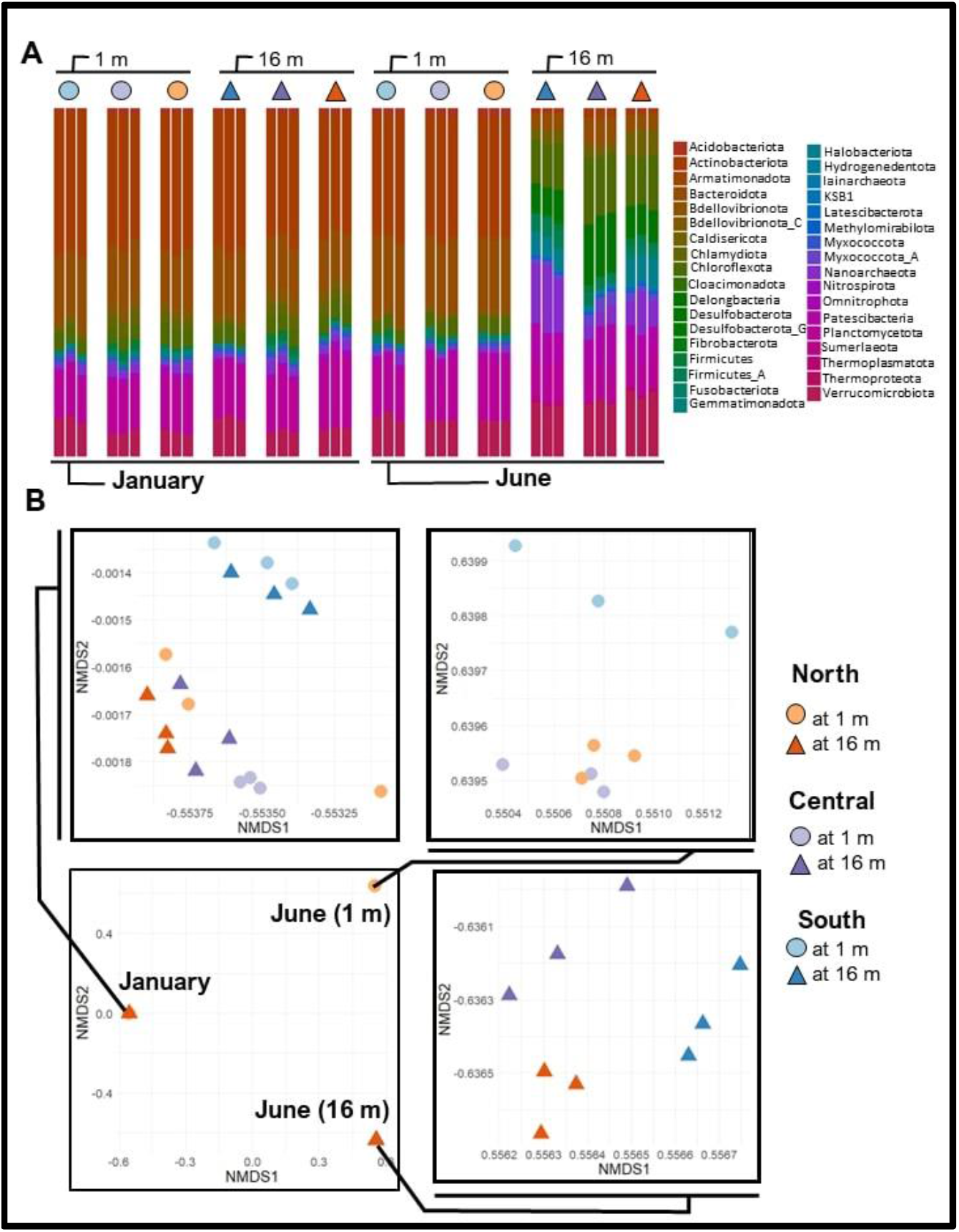
**(A)** Diversity of phyla among samples, with dominant phyla (*Cyanobacteria* and *Proteobacteria*) removed to more clearly reflect community differences between depths and across seasons. **(B)** Due to highly diverse community assemblages, ordination analysis of membership has been separated into January samples, June surface samples and June hypolimnetic samples.

While differences between depths in genome taxonomy were evident in June at the phyla level (Fig. 3A), differences in genome taxonomy between the June surface and January (both depths) communities were structured at lower levels of classification (Fig. S2). Given the diversity of lineages, we used a nonmetric multidimensional scaling (NMDS) analysis of MAG relative expression within the transcriptome (hereafter referred to as genomes) to qualitatively describe differences in microbial membership within Lake Yojoa in space and time. Due to the dissimilarity of genomes among samples, differentiation among all dates and depths within a single two-dimensional ordination analysis was not possible (Fig. 3B).

Therefore, to more clearly describe differences in microbiome composition across seasons and among sampling stations, we performed separate ordination analyses on January (both depths), June (surface), and June (hypolimnion) samples (Fig. 3B). In January, we saw no separation in ordination space by depth, as would be expected when the water column was mixed. However, genomes in southern sampling station (at both depths) differed from the other two sampling locations in January. Similarly, the June surface genomes at the southern sampling point were also distinct from the other two locations, which were more similar to each other. In the hypolimnion in June, genomes at the three sampling locations were all dissimilar (Fig. 3B), likely reflective of decreased physical mixing in the hypolimnion during stratification and station specific geochemical differences (Fig. 2).

To specifically address contribution of the microbiome to N cycling, we also identified the taxa of the genomes that expressed N cycling genes of interest (Table S2). In January, all taxa expressing N cycling genes of interest were present at both depths (Fig. S3). Conversely in June, lineages with N metabolism genes of interest were much more distinct between depths. Many of the identified N cycling families were present only in the hypolimnion in June, while a smaller proportion were present at both 1m and 16 m (and no lineages appeared only at 1 m) (Fig. S3). *Dominant microbial N metabolisms*

To characterize the dominant N metabolic pathways in our samples, we calculated relative expression (i.e., number of transcripts) of genes of interest (Table S2) for each station on each date, at each depth. Relative expression within triplicates was then summed within each date and depth hereafter referred to as treatments (i.e., June at 1 m, June at 16 m, January at 1 m, and January at 16 m). We then compared gene expression across each of the four treatments. This allowed us to identify which genes were upregulated or downregulated for a particular treatment relative to all other treatments.

Consistent with trends in the geochemistry data and microbiome membership, we observed similar gene expression for genes involved in N metabolism in January between depths and across stations and dissimilarities in N metabolism gene expression between depths in June (Fig. 4A). In January, at both 1 m and 16 m, respiratory NO_2_^-^ reductases (*nirB, nirD, nrfA, nrfH*), NO-forming NO_2_^-^ reductase (*nirK*), and assimilatory NO_3_^-^ reductases (*narB, nasA, nirA*) were the most highly expressed N metabolism genes. Whereas these three gene groups were similarly expressed at both 1m and 16 m, assimilatory NO_3_^-^ reductases were upregulated relative to other NO_3_^-^/NO_2_^-^ reduction pathways at 1 m compared to at 16 m (Fig. 4). Despite oxic conditions throughout the water column during the January sampling event, we also observed expression of genes involved in several anaerobic metabolisms in January (Fig. 4D-E), including those associated with denitrification beyond NO formation (nitric oxide reductase, *norB/C*, and nitrous oxide reductase, *nosZ*, which was downregulated relative to proceeding steps in denitrification).

**Figure 4.**
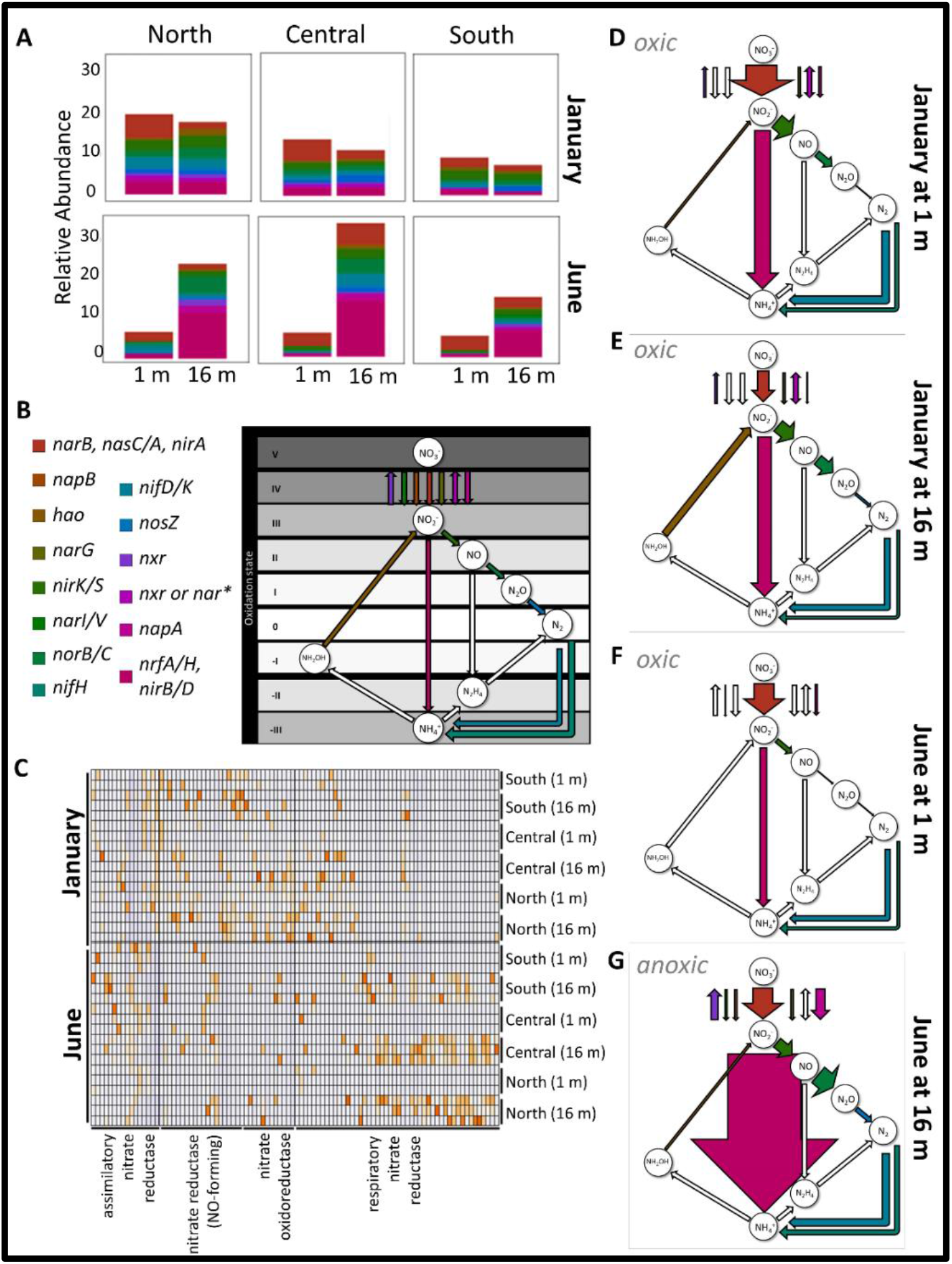
**(A)** N metabolism gene expression (relative abundance across all samples with summed triplicates), grouped by function, across locations, depths and season. **(B)** N cycling pathways assessed in panels D-G, putative transformations. *Differentiation between *nar* and *nxr* expression was not always possible. **(C)** Heatmap of gene expression of nitrite transformation pathways. **(D-G)** Conceptual diagram of dominant N transformation pathways across seasons and 1 m and 16 m, shown in arrows weighted by expression. Relative abundance data from panel A has been summed by treatment group. Unfilled arrows represent the complete absence of gene expression.

The most pronounced differences we observed in N cycling associated gene expression occurred between the oxic surface and the anoxic hypolimnion in June (Fig. 4C). In the surface, dominant N metabolisms mirrored those in January at 1 m with assimilatory NO_3_^-^ reductases being most highly expressed followed by respiratory NO_2_^-^ reductases and NO-forming NO_2_^-^ reductase. Also, similar to January, *nosZ* in the June surface was downregulated relative to previous steps in denitrification. In the hypolimnion, respiratory NO_2_^-^ reductases were overexpressed relative to all other gene categories (Fig. 4). Within the respiratory NO_2_^-^ reductases, *nrfA* was the most highly expressed gene followed by *nirB, nrfH* and *nirD*, respectively. Collectively, these genes were the most highly expressed across all treatments and stations but had the largest differential enrichment in the June hypolimnion, indicating strong selection pressure for the pathways these genes are involved in. Using weighted arrows to reflect combined respiratory nitrate reduction (*nrfA/H* and *nirB/D*), we demonstrate pronounced enrichment of DNRA pathways in June hypolimnion relative to other strata and dates (Fig 4).

Due to the notable upregulation of respiratory NO_2_^-^ reduction genes, we identified the taxa assigned to all genomes that expressed *nirB, nirD, nrfA*, and *nrfH*. In addition to lineages undefined at the order level, we found 18 orders that were responsible for expressing respiratory NO_2_^-^ reductases (Fig. S4). Order *Desulfomonilales* was the most abundant lineage followed by *Anaerolineales, Methylococcales*, and an order of *Myxococcota* (*UBA796*). Looking only at lineages expressing *nrfA*, the gene most commonly associated with DNRA (51), 11 of the initially identified 18 lineages (which were performing respiratory NO_2_^-^ reduction) were putatively performing DNRA in the hypolimnion (Fig. S5). Lineages from families *Anaerolineaceae, Desulfobulbaceae*, and *JAFGLY01* were the most abundant transcribers of *nrfA* with expression being significantly highest in *Anaerolineales Anaerolineaceae*.

### Microbial organic N mineralization

To examine additional potential microbial mechanisms that may be responsible for the previously described (17) accumulation of hypolimnetic NH_4_^+^, we profiled metatranscriptomic data for expressed peptidases (genes that mineralize organic N into amino acids), organic N transporters (genes for cellular uptake of smaller organic N compounds), amino acid transformers (genes for mineralization of amino acids leading to NH_3_^+^), and ureases (genes that hydrolyze urea to produce NH_3_^+^) (Fig. 5). Collectively, more expression of organic N genes occurred in June relative to January with the most occurring in the hypolimnion, suggesting that mineralization of organic N may also play a role in the previously observed increase in hypolimnetic NH_4_^+^ during stratification (Fig. 2E) (17).

**Figure 5.**
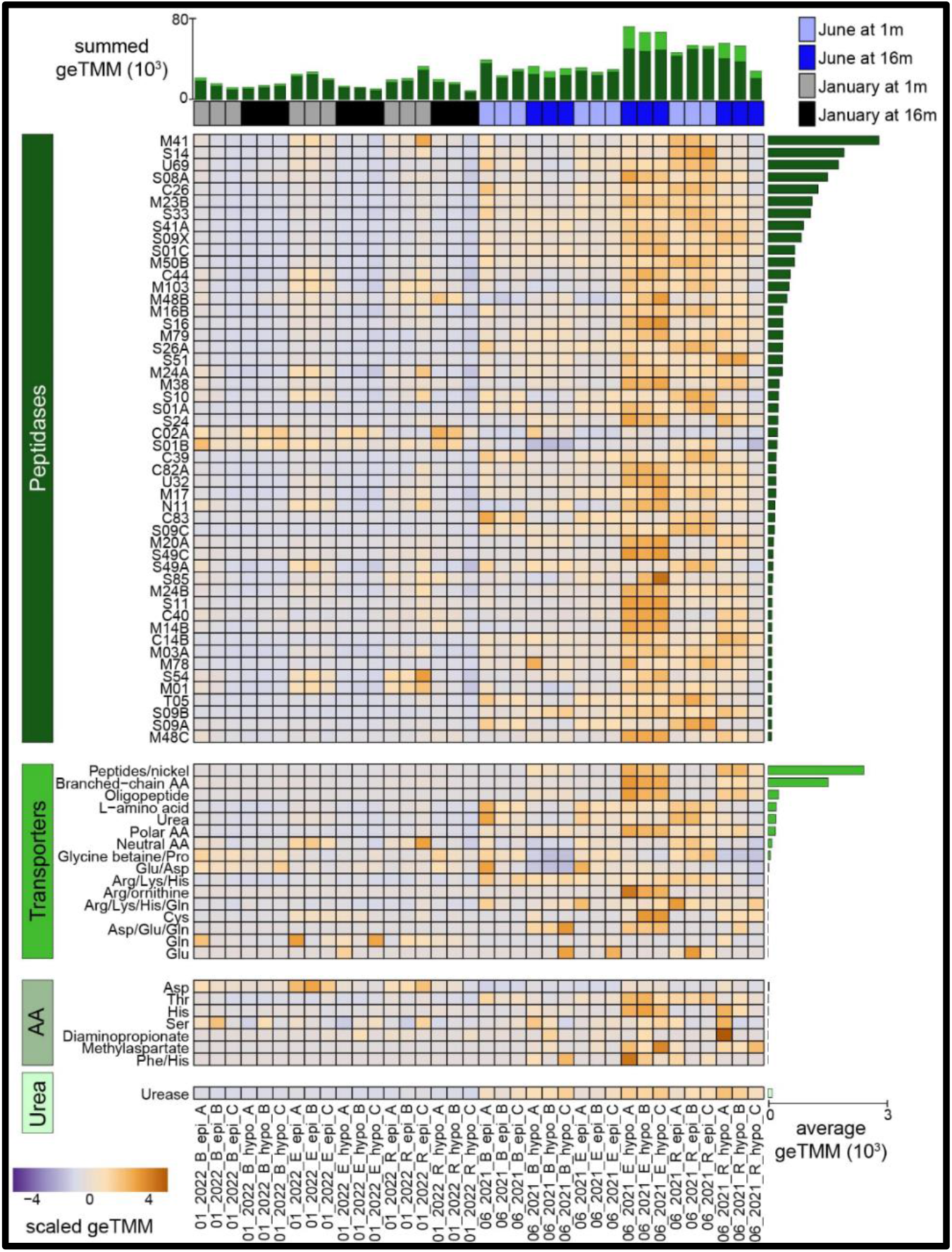
Organic nitrogen mineralization expression in Lake Yojoa. Heatmap shows the expression of peptidases (at the family level, top 50 shown), transporters, amino acid transformations, and urease. Expression is scaled within row and ordered by average transcription across all samples within each category, as depicted by side bar plot. Top stacked bar plot shows the summed expression within each sample, with bars colored by category of organic nitrogen shown on the left side of heatmap.

Categorization of these genes revealed expression in all categories of organic N utilization, including 37,004 peptidases, 5,403 organic N transporters, 214 amino acid transforming genes, and 97 ureases. Genes within the M41 and S14 families had the most expression, with M41 being made up of exopeptidases (cleaves peptide bonds at a terminal end of a protein or peptide) and S14 being made up of endopeptidases (cleaves peptide bonds of non-terminal amino acids). Highly expressed extracellular families included S08A and M23B, made up of endopeptidases that have the potential to cleave after hydrophobic residues and lyse bacterial cell wall peptidoglycans, respectively. Nearly all lineages contributed to peptidase expression, with members of *Cyanobacteria, Chloroflexota*, and *Proteobacteria* phyla being the most highly expressed.

Similarly, *Cyanobacteria* and *Chloroflexota* phyla had the highest average expression within the transporter category. *Cyanobacteria* also had the most expression in the two most expressed NH_4_^+^ forming reactions (urease and aspartate transformations). These results highlight members of the *Cyanobacteria* as key contributors to the N cycle in Lake Yojoa, with the highest average expression in all categories genes involved in organic N cycling.

## Discussion

In the Lake Yojoa microbiome, membership and N metabolic gene expression appeared to be primarily influenced by Lake Yojoa’s monomictic stratification regime that resulted in pronounced redox differences between water column strata and seasonal depth specific changes in electron donor and acceptor availability. While there was minimal difference in microbiome membership between depths during January when the water column was mixed, there were distinct differences in membership and gene expression between the surface and the hypolimnion in June. Here we explore these seasonal and depth discrete differences in Lake Yojoa’s microbiome and the role it plays in N cycling by discussing the key results from analyses of the oxic water samples (January at both 1 and 16 m and June at 1 m) within the context of previously observed intra-annual dynamics of the Lake Yojoa ecosystem. We then describe the observed spatial and temporal trends in organic N mineralization across seasons and depths. We conclude the discussion by focusing on the observed NO_3_^-^ and NO_2_^-^ reduction pathways we observed in the anoxic June hypolimnion and characterizing and assessing the putative role of DNRA in NH_4_^+^ accumulation within Lake Yojoa’s warm anoxic water hypolimnion.

In June, the downregulation of *napA* (periplasmic nitrate reductase) in the surface (relative to January) was likely due to low concentrations of NO_3_^-^, which is depleted in surface waters by June (17). This low abundance of inorganic N in the photic zone in June relative to January is consistent with previously described N and P colimitation in June but P limitation in January which allows inorganic N to persist at higher concentrations during the mixed water column months (17). As with NO_3_^-^, NH_4_^+^ was low in the June surface. The absence of *hao* (hydroxylamine oxidoreductase) expression is consistent with expectations of low nitrification potential (given low NH_4_^+^) availability at this time. Conversely, measurable *hao* expression in January was consistent with the increased NH_4_^+^ concentration we observed relative to June surface samples. Despite measurable quantities of NH_4_^+^ at all sampling points, we were unable to identify expression of ammonia monooxygenase (*amoA*) for any of the treatments. We assessed our unbinned assembly fractions for *amo*, a function missing in our MAG database. From our assemblies, we recovered a single copy of an *amoA* on a scaffold (<5000 bp) likely assigned to an unbinned *Nitrosomonas*.

In addition to the expected aerobic metabolisms, we also identified gene expression of putatively anaerobic pathways associated denitrification and DNRA (e.g., *nirK/B, norB/C, nosZ, nrfA*) in the oxic water column. One explanation of this observation is the presence of aerobic denitrifiers as have been identified in other aquatic ecosystems which experience frequent fluctuation between oxic and anoxic conditions (26). However, expression of genes associated with anaerobic metabolism is more likely explained by the presence of anoxic microsites within the water column (e.g., on sinking particles or other colonized aggregates). Anerobic metabolisms have been demonstrated to be significant contributors to ecosystem scale biogeochemical cycling in bulk oxic environments (27) and sinking particulates are hotspots of anaerobic metabolism in both marine and lake ecosystems (28-30). The presence of anoxic microsites also explains expression within genomes identified as strict anaerobes, such as *Desulfomonilia*, in the June surface waters and January water column. In Lake Yojoa, sinking particulates (from primary productivity and fish waste associated with aquaculture) provide ample substrate for the formation of anoxic microsites.

Expression of genes related to organic N mineralization pathways was typically higher in June samples compared to January for most peptidases, organic N transporters, amino acid transformers, and ureases. Notable exceptions to this trend include two peptidases, C02A (cysteine) and S01B (Serine), and one amino acid transformer, aspartic acid which were upregulated in January relative to June. For all other peptidases, expression was greatest in June, particularly at 16 m in the central sampling location, nearest to the fish pens (Fig. 1). Mineralization genes were also upregulated at 1 m and, to a lesser extent, 16 m in the northern most sampling point. These mineralization hotspots likely reflect Lake Yojoa’s dominant watershed derived nutrient sources (three of the six major tributaries are located in the northwest basin of the lake) and industrial aquaculture, also located in the north central basin (Fig. 1). As dominant wind direction blows north to south, surface particles from the aquaculture operation are transported south to the central location, likely supplying the hypolimnion in mineralizable N-rich organic matter. The outsized role the aquaculture plays in the nutrient budget of Lake Yojoa suggest that mineralization of fish waste may be a principal driver and perhaps the most parsimonious explanation for the previously observed accumulation of hypolimnetic NH_4_^+^.

In the June hypolimnion, we saw an upregulation in *nrfA* relative to June surface samples. This distinct difference in gene expression between the top and bottom strata mirrors *nrfA* patterns previously reported in Lake Alchichica, Mexico (15) where N-associated gene abundances was also driven by seasonal patterns in stratification. However, gene expression is imperfectly correlated to protein expression (31, 32) and rarely, with some notable exceptions, correlates with rate processes (33). This limits the biogeochemical inference that can be drawn between the relative abundance of transcripts and the role of DNRA and other pathways which compete for NO_3_^-^ (e.g., denitrification). Therefore, in the absence of direct rate measurements, we are unable to determine the relative proportion of NO_2_^-^ reduced by DNRA versus denitrification. However, we can conclude that in the June hypolimnion, across all locations, DNRA pathway genes (*nrfA/H* and *nirB/D*) were enriched relative to denitrification pathway genes (*nirK, nirS, norB*, and *nosZ*) and a principal gene associated with the anammox pathway (*hzsA*) was absent. Furthermore, *nrfA* expression was negatively correlated to NO_3_^-^ concentrations in the June hypolimnion (R^2^= 0.84, *p*< 0.005). This suggests that DNRA is increasingly competitive under conditions of low NO_3_^-^ availability (34) and is consistent with the observed annual NH_4_^+^ accumulation that occurs in Lake Yojoa.

The upregulation of *nrfA* for all three stations in Lake Yojoa, which are kilometers apart, differ in maximum depth, and are different distances from large nutrient sources, suggests DNRA in the hypolimnion of Lake Yojoa is a ubiquitous and perhaps a key mechanism for NH_4_^+^ accumulations. Further supporting the important role of DNRA in Lake Yojoa’s N cycle is the presence of several *nrfA* expressing lineages (such as *Anaerolineales, Burkholderiales*, and *Desulfobulbales*) that have been identified as performing DNRA in other ecosystems (Fig. S5) (35-37). The majority of these lineages were absent in the oxic surface in June, though *nrfA* gene expression in a subset of those orders (i.e., *Burkholderiales, Phycisphaerales, Tepidisphaerales* and *UBA1135*), was present at both depths, likely supported by the anoxic microsites discussed above.

Our study highlights the putative contributions of mineralization and DNRA to the hypolimnetic NH_4_^+^ pool of a large tropical lake. Although we acknowledge that multiple other pathways may play an important role in the accumulation of hypolimnetic N (e.g., sediment derived NH_4_^+^ flux (22), mineralization of fish waste or interactive effects of Lake Yojoa’s virome on N cycling (38, 39)), DNRA, because of its competition with inorganic N ecosystem loss pathways, remains a critical pathway for assessing annual N dynamics, particularly in systems, like Lake Yojoa, that experience seasonal N limitation (17). Determining controls on competing NO_3_^-^/NO_2_^-^ reduction pathways the warm anoxic waters in Lake Yojoa and other low-latitude lakes, are critical steps in broadening our mechanistic understanding of microbially driven biogeochemical processes that influence the trophic state of tropical freshwater ecosystems. Only by establishing this fundamental ecological knowledge can we hope to understand why and how tropical lakes are responding to a changing world.

## Conclusion

By identifying the dominant N metabolisms which govern intra-annually variable reactive N availability in Lake Yojoa, we provide new insights into the microbial pathways of these understudied warm, seasonally anoxic ecosystems. Descriptions of such pathways may contain clues that distinguish tropical lake biogeochemistry from temperate lake biogeochemistry. Our results highlight the degree to which the largely undescribed taxonomic and functional diversity in such ecosystems define ecosystem scale nutrient fluxes. We have also demonstrated the need to define controls and constraints on DNRA, in addition to mineralization. By better understanding the microbial assemblages and emergent metabolisms in tropical lakes, particularly in hypolimnions, we may begin to understand how lakes, like Lake Yojoa, as well as lakes at higher latitudes under future climate scenarios, function under contemporary and eminent environmental stressors.

## Methods

### Field Sampling

In June of 2021 and January of 2022, we collected water for nutrient analyses (NH_4_^+^, NO_3_^-^, total phosphorus, dissolved organic carbon) from three stations within the lake (Fig. 1) at 1 and 16 m depth using an opaque Van Dorn water sampler. Stations were chosen to be approximately equidistant from each other to capture potential spatial heterogeneity of the pelagic zone of Lake Yojoa. Methods for samples collection, preservation, transportation, and analysis as well as thermophysical profile measurements are available in previously published works on Lake Yojoa (17, 22).

For DNA and RNA sample collection, 100-300 ml water samples were concentrated onto 0.22-μm pore size Sterivex filters (Sigma Aldrich Cat. # SVGP01050) using sterile 60 ml luer-locking syringes. Water was passed through filters until filters were at capacity. Filters were then quickly purged with air to remove excess water and 3 ml of RNALater (Sigma Aldrich Cat. # R0901500ML) was added. Sealed Sterivex filters were then placed in individual whirl-paks and placed in a cooler.

Although we only compared two sampling dates, the contrast between June 2021, when stratification was fully developed, juxtaposed against January 2022, when the water column was fully mixed, is representative of the geochemical conditions of the two phases of stratification that Lake Yojoa experiences. No disruption to stratification was identified prior to June sampling (22) and lake microbial communities have been shown to be resilient (on the scale of days) to water column mixing disturbance (40). Thus, the samples taken in June and January are likely representative of the stratified and mixed water column seasons, respectively.

### Extraction

Samples for metagenomics were taken during both sampling events (June 2021 and January 2022) from each location (North, Central and South) and each depth (1 m and 16 m) and extracted for DNA (n=12 metagenomes). To maximize genome recovery and build a robust metagenomic assembled genome (MAG) database for metatranscriptome mapping, we performed additional metagenomic sequencing (n=15 metagenomes) on samples collected during 2019 and 2020 sampling campaigns (17, 22). Samples for RNA were paired to the June 2021 and January 2022 metagenomic sampling and collected from the same locations and depths in triplicate (n=36 metatranscriptomes). DNA and RNA were coextracted using ZymoBIOMICS DNA/RNA Miniprep Kit (Zymo Research Cat. # R2002) coupled with RNA Clean & Concentrator-5 (Zymo Research Cat. # R1013). Samples were eluted in 40 μL and stored at −20 °C until they were sent for sequencing as described below.

### Metagenomic assembly, binning, and annotation

Genomic DNA was prepared for metagenomic sequencing using Plate-based DNA library preparation on the PerkinElmer Sciclone NGS robotic liquid handling system at the Joint Genome Institute. Briefly, one nanogram of DNA was fragmented and adapter ligated using the Nextera XT kit (Illumina) and unique 8bp dual-index adapters (IDT, custom design). The ligated DNA fragments were enriched with 12 cycles of PCR and purified using Coastal Genomics Ranger high throughput agarose gel electrophoresis size selection to 450-600bp. The prepared libraries were sequenced using an Illumina NovaSeq following a 2x150nt indexed run recipe.

Resulting fastq files were assembled and binned using the GROWdb pipelines (https://github.com/jmikayla1991/Genome-Resolved-Open-Watersheds-database-GROWdb/tree/main/Yojoa_Honduras_Lake). Briefly, three assemblies were performed on each set of fastq files and binned separately: (1) Read trimming with sickle (v1.33), assembly with megahit (v1.2.9), and binning with metabat2 (2.12.1) (2) Read trimming with sickle (v1.33), random filtering to 25% of reads, assembly with idba-ud (1.1.0), and binning with metabat2 (2.12.1) (3) Bins derived from the JGI-IMG pipeline were downloaded. All resulting bins were assessed for quality using checkM (v1.1.2) and medium and high-quality MAGs with >50% completion and <10% contamination were retained.

For paired June 2021 and January 2022 metagenomes, subassemblies were also performed. Specifically trimmed reads from 12 samples were individually mapped to medium and high-quality MAGs derived from the three assembly types described above using bbmap (perfectmode=t) (41). Unmapped reads for each sample were then assembled with idba-ud (1.1.0) (42) and binned with metabat2 (2.12.1) (43). These bins were also assessed for quality using checkM (v1.1.2) (44) and MAGs with >50% completion and <10% contamination were retained in the database. The resulting 1,771 MAGs across all samples and assemblies were dereplicated at 99% identity using dRep (v2.6.2) (45) to obtain the dereplicated Yojoa MAG database (n=572 MAGs). MAG taxonomy was assigned using GTDB-tk (v2.0.0) (46) and annotated using DRAM (47). Methods for classifying gene homologs provided in Supplementary Text 1.

### Metatranscriptomic mapping and analysis

RNA was prepared from metatranscriptome sequencing according to JGI established protocols. A summary of JGI’s protocols is available in Supplementary Text 2. Resulting fastq files were mapped via Bowtie2 (-D 10 -R 2 -N 1 -L 22 -i S,0,2.50) (48) to the dereplicated Yojoa MAG database (n=572 MAGs). Sam files were transformed to bam files using samtools filtered to 97% id using reformat.sh (49) and name sorted using SAM Tools (50). Transcripts were counted each gene with featureCounts (51). Counts were transformed to geTMM in R using edgeR package (52).

### Statistical analysis and figure generation

Statistical analyses and figure generation were performed in R (version 4.1.2). Significant differences in geochemical and thermophysical structure data between depths and/or seasons were assessed using one-away analysis of variance (ANOVA). Circular dendrograms were created using RAWgraphs (53).

## Acknowledgements

Samples were sequenced and processed as a part of the Genome Resolved Open Watersheds (GROW) effort to sequence global watersheds. We thank Tyson Claffey and Richard Wolfe for Colorado State University server management. Additionally, we thank Juan Carlos Sorto for being instrumental in sample collection. This manuscript was improved by feedback from Josué Rodríguez-Ramos.

## Data availability statement

Raw reads and metagenome assembled genomes are publicly available on NCBI under Bioproject PRJNA946291. Dereplicated metagenome assembled genomes are also publicly available on Zenodo (https://zenodo.org/record/8035120) along with annotations, quality statistics, mapping tables, and geochemistry and thermophysical profile data.

