## Supplementary Tables and Figures for "Dominant nitrogen metabolisms of a warm, seasonally anoxic freshwater ecosystem revealed using genome resolved metatranscriptomics"

**Table S1.** Mean ± standard error of geochemical parameters across the three sampling locations.

| | **$NH_4^+$ (μM)** | **$NO_3^-$ (μM)** | **DOC (μM)** | **TP (μM)** |
|---|---|---|---|---|
| January (1 m) | 38.01 ± 3.19 | 15.45 ± 0.25 | 200.75 ± 7.21 | 0.56 ± 0.03 |
| January (16 m) | 39.86 ± 3.38 | 15.50 ± 0.27 | 208.75 ± 7.73 | 0.60 ± 0.21 |
| June (1 m) | 3.85 ± 0.67 | 2.04 ± 0.43 | 225.26 ± 25.91 | 1.23 ± 0.40 |
| June (16 m) | 27.25 ± 5.18 | 1.79 ± 0.21 | 154.87 ± 14.86 | 0.83 ± 0.28 |

**Table S2.** Nitrogen metabolism genes considered in this analysis.

| **Kegg ID** | **Gene** | **Description** |
|---|---|---|
| K00367 | *narB* | assimilatory nitrate reductase |
| K00372 | *nasC, nasA* | assimilatory nitrate reductase |
| K00366 | *nirA* | assimilatory nitrite reductase |
| K02568 | *napB* | Nitrate reductase, cytochrome c-type protein |
| K10535 | *hao* | hydroxylamine oxidase |
| K00368 | *nirK* | nitrite reductase (NO-forming) |
| K15864 | *nirS* | nitrite reductase (NO-forming) |
| K00374 | *narI, narV* | nitrate reductase 1 |
| K04561 | *norB* | nitric oxide reductase |
| K02305 | *norC* | nitric oxide reductase |
| K02588 | *nifH* | nitrogenase iron protein |
| K02586 | *nifD* | nitrogenase molybdenum-iron protein |
| K02591 | *nifK* | nitrogenase molybdenum-iron protein |
| K00376 | *nosZ* | nitrous-oxide reductase |
| K00371 | *narH, narY, nxrB* | nitrite oxidoreductase |
| K00370 | *narG, narZ, nxrA* | nitrite oxidoreductase |
| K02567 | *napA* | periplasmic nitrate reductase |
| K00362 | *nirB* | respiratory nitrite reductase |
| K00363 | *nirD* | respiratory nitrite reductase |
| K03385 | *nrfA* | respiratory nitrite reductase |
| K15876 | *nrfH* | respiratory nitrite reductase |

### Supplementary Figures

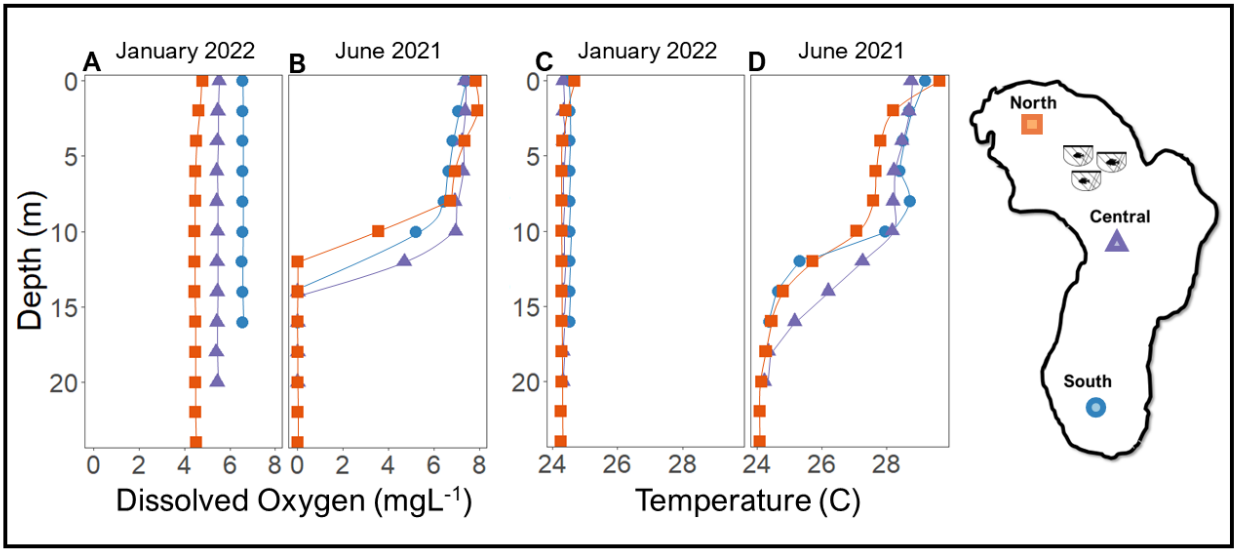


**Figure S1.** Thermophysical structure in Lake Yojoa. **(A)** Dissolved oxygen at 2 m intervals in January 2022 and June 2021**. (B)** Temperature at 2 m intervals in January 2022 and June 2021.

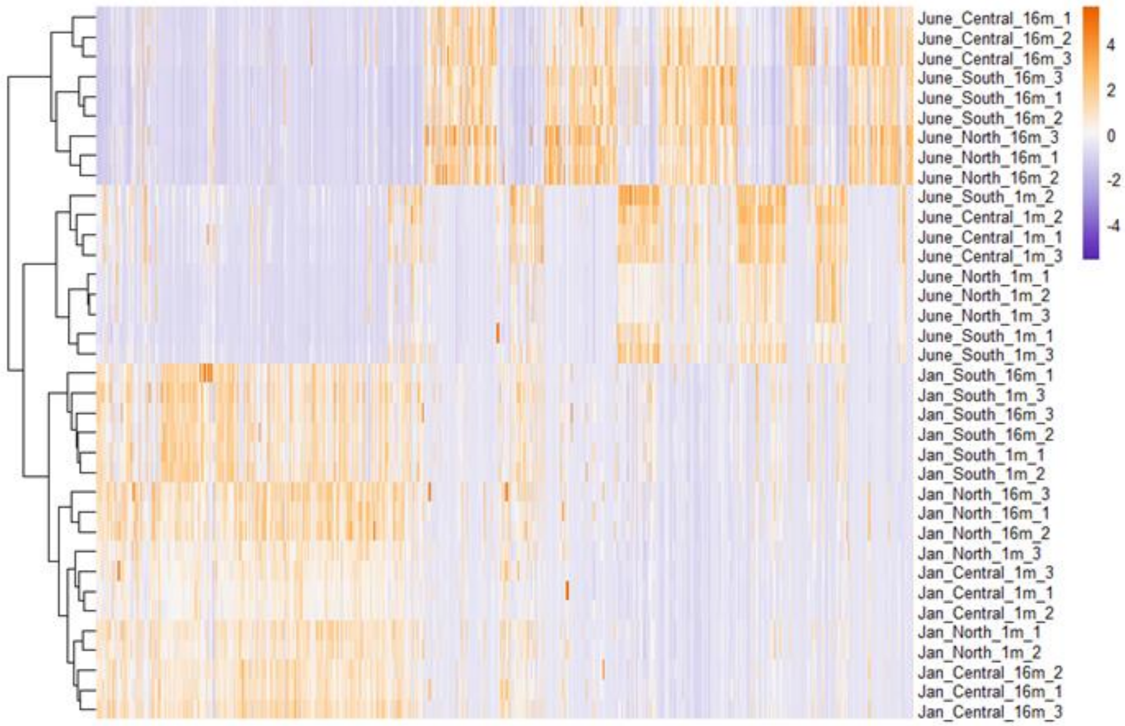


**Figure S2.** MAG relative expression heatmap with hierarchical clustering.

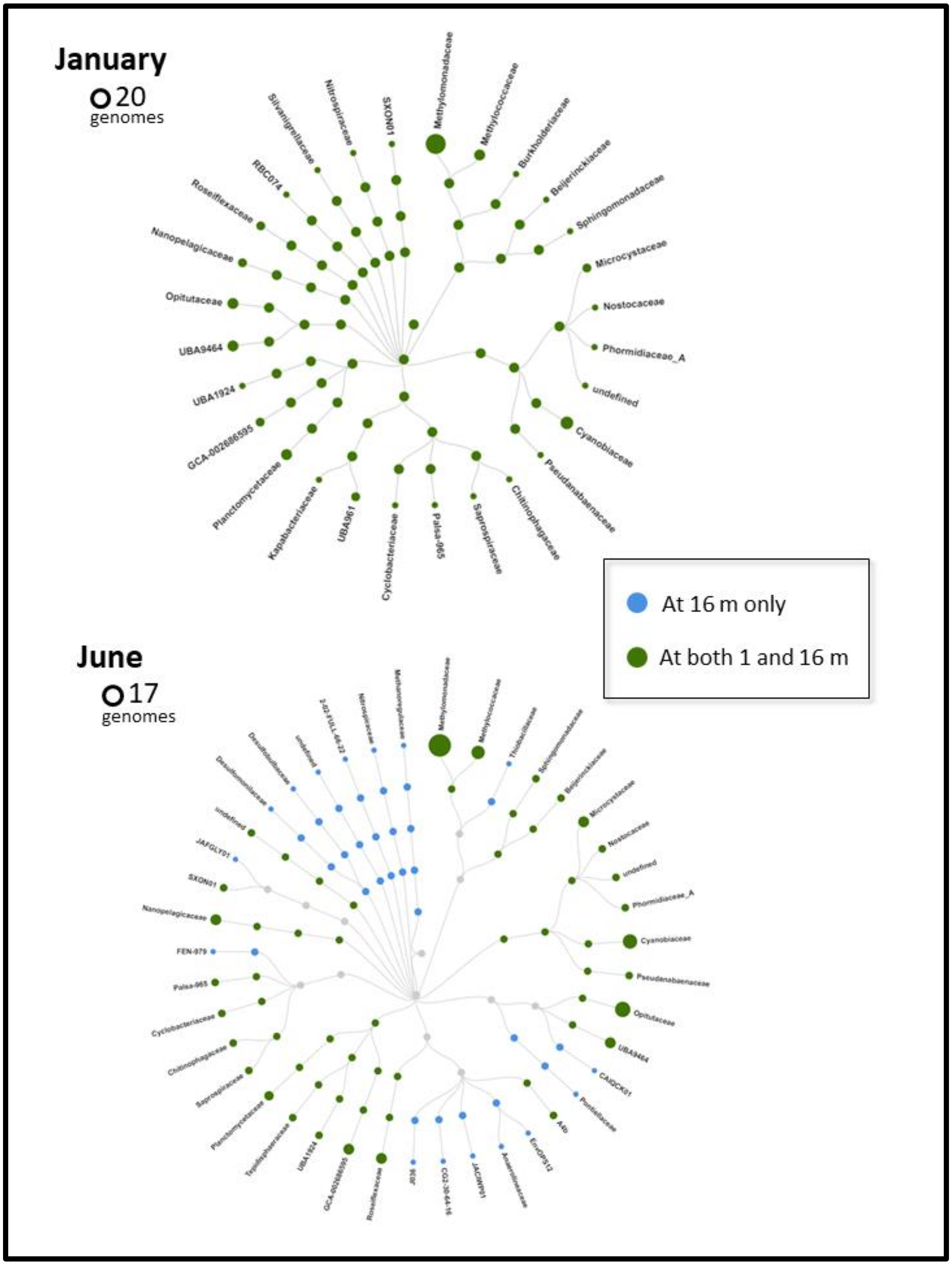


**Figure S3.** Taxonomic level (Domain, Phylum, Class, Order, Family) assigned by GTDB-tk to genomes expressing N metabolism genes of interest (Table S1) in January (n= 20) and June (n=17).

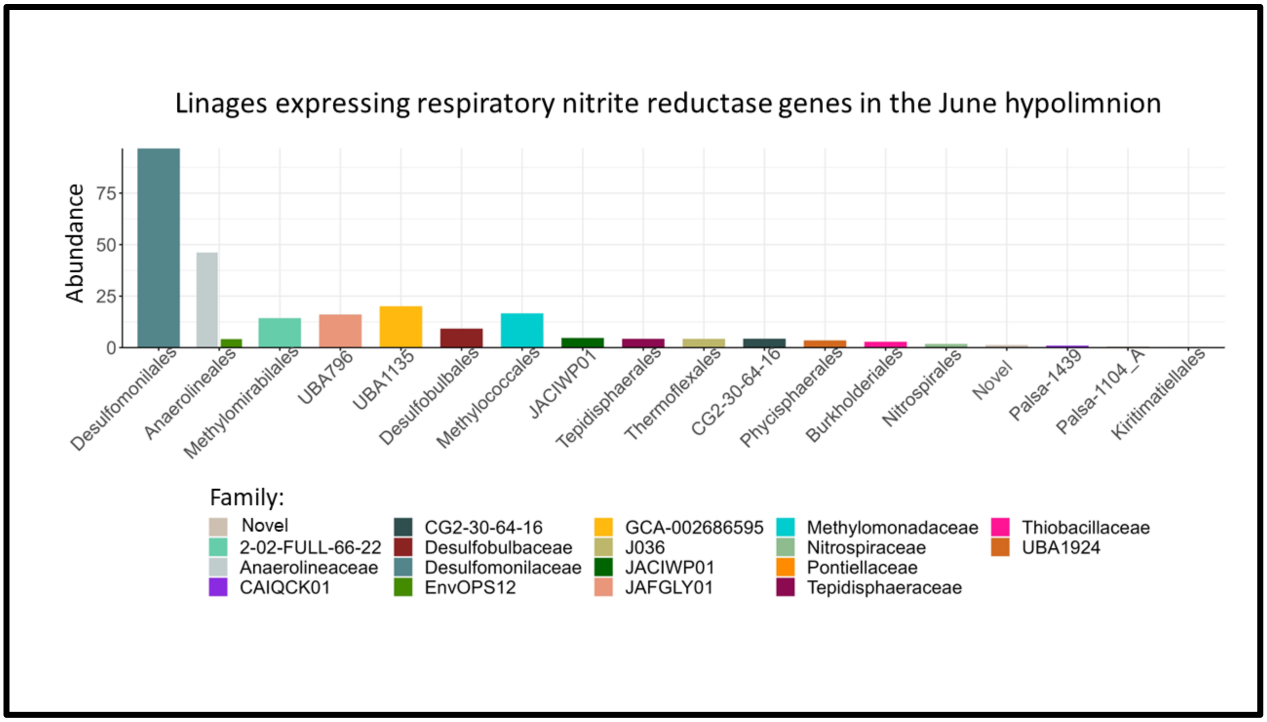


**Figure S4.** Lineages of MAGs expresing genes related to respiratory nitrite reduction (Table S1). Abundance refers to total expression.

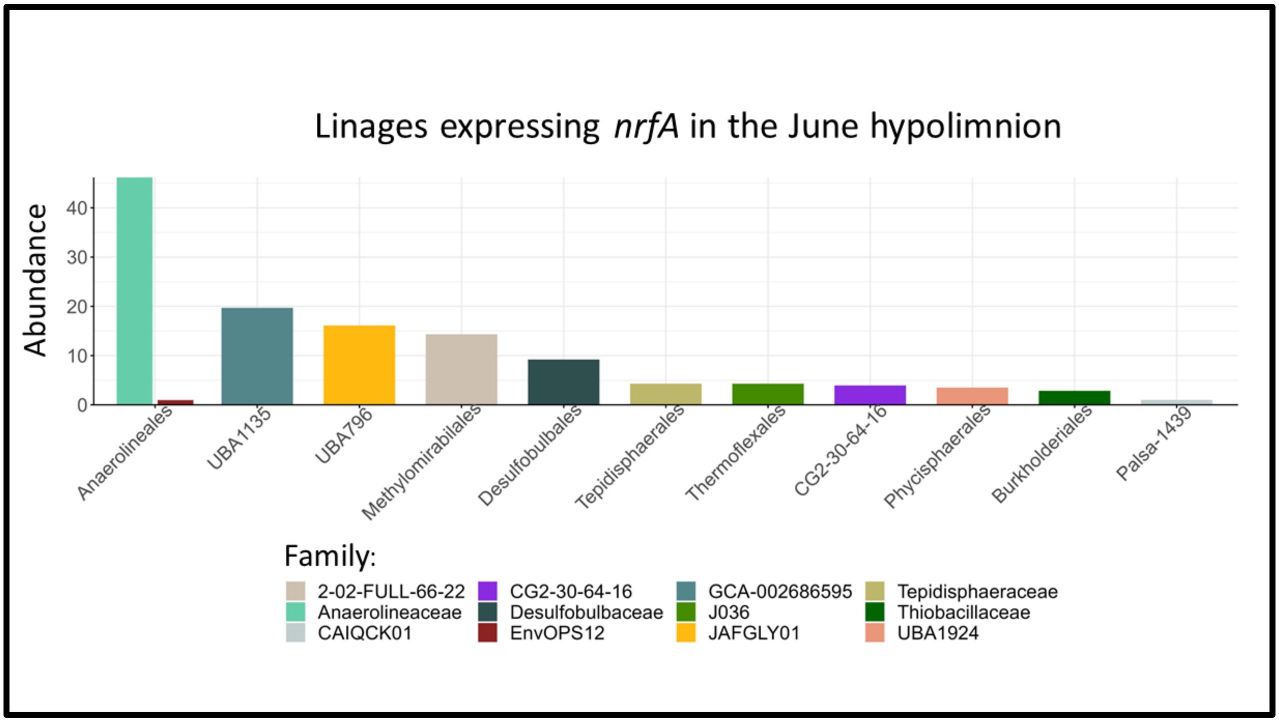


**Figure S5.** Lineage of MAGs expressing *nrfA* in the June hypolimnion. Abundance refers to total expression.
