## Supplementary Text 1 for "Dominant nitrogen metabolisms of a warm, seasonally anoxic freshwater ecosystem revealed using genome resolved metatranscriptomics"

**Classifying homologs in the MAG database**

Additional phylogenetic analyses were performed on genes annotated as respiratory nitrate reductase (*nar*), nitrite oxidoreductase (*nxr*), ammonia monooxygenase (*amo*), or methane monooxygenase (*pmo*) to better assign the role of MAGs derived from Yojoa in specific N cycling pathways. To do this 100 *nxr*/*nar* and 65 *pmo*/*amo* reference sequences were downloaded(1-4). Each set of reference sequences were combined with amino acid sequences of homologs from the Yojoa MAG database, aligned separately using MUSCLE (v3.8.31), and run through an in-house script for generating phylogenetic trees (https://github.com/WrightonLabCSU). This resulted in two phylogenies, one for *nxr*/*nar* and one for *pmo*/*amo*, which were used to classify gene homologs in the MAG database.

*pmoA/amoA*

No putative amoA. All hit to methane (pmoA) or alkane.

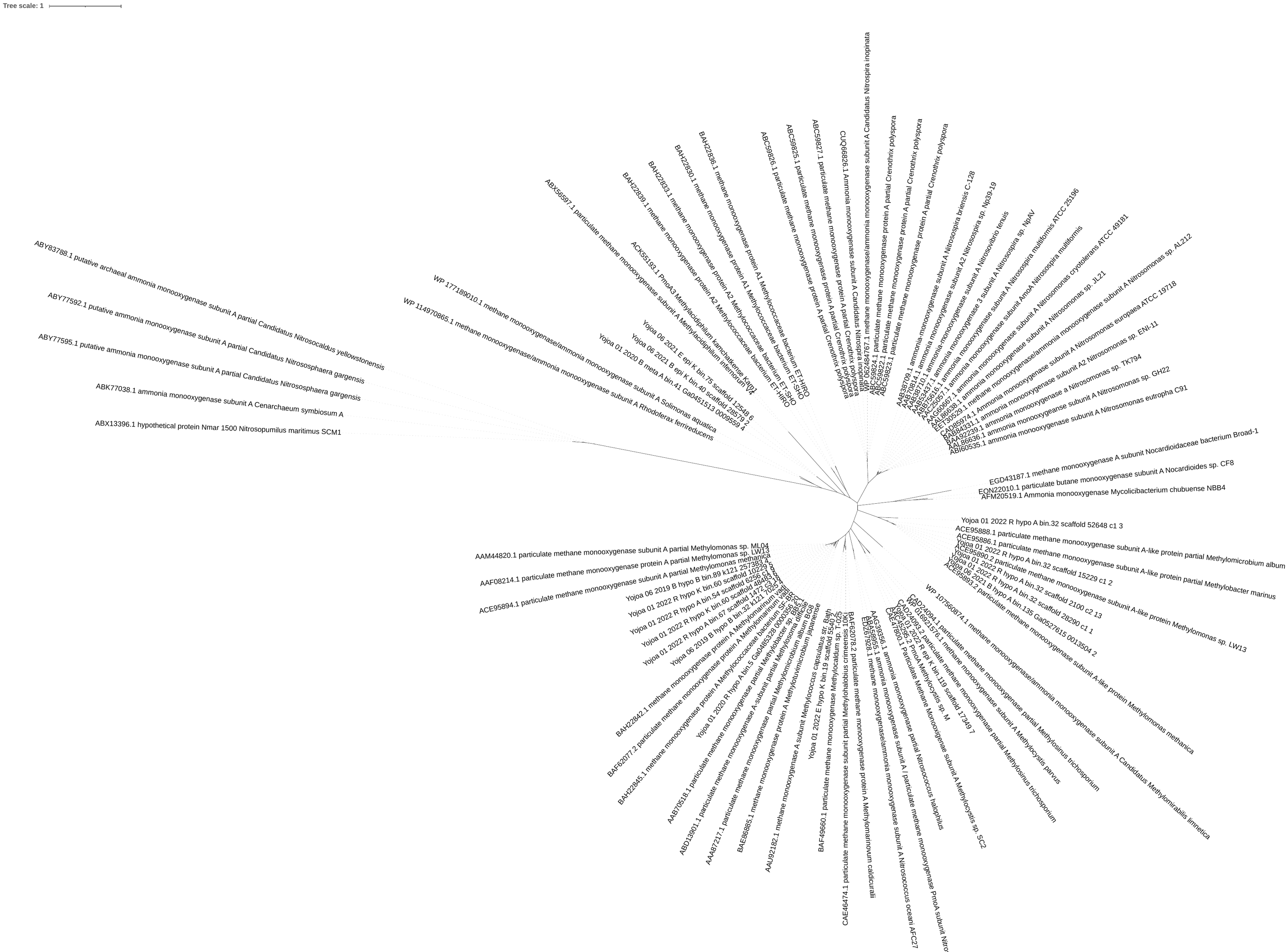

Tree from dereplicated MAGs.

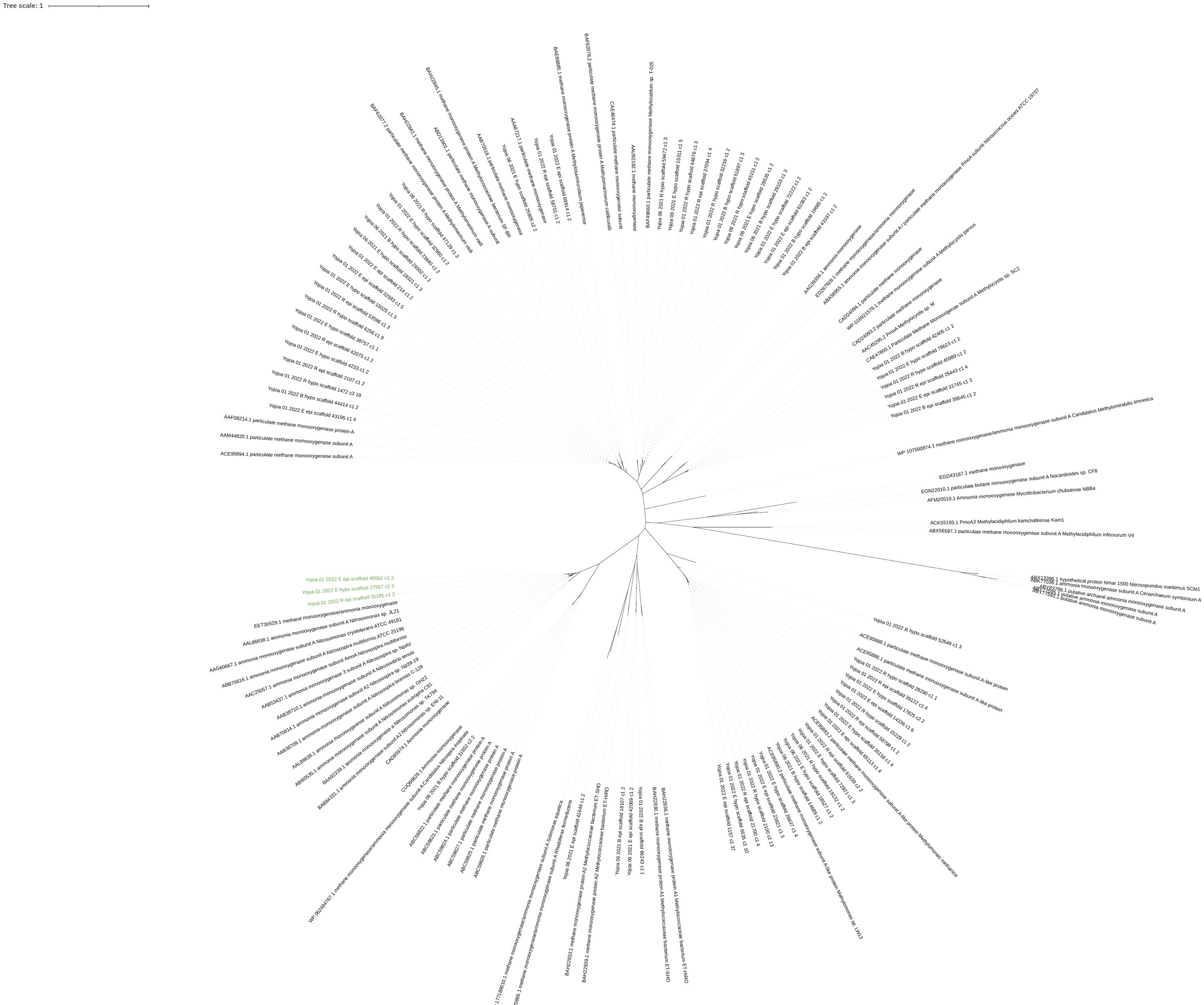

Tree from assemblies

Putative *amoA* from assemblies:

>Yojoa_01_2022_R_epi_scaffold_31185_c1_2

MSRTDEILAAAKMPPEAVRMSRYIDAVYFPILCILLVGTFHMHFMLLAGDWDFWLDWKDRQWWPVVTPIVGVMYCAALQYYLWVNYRLSYGATLCIVCLLVGEWLTRYWGFYWWSHYPLNFVLPSTMIPGALMLDTILLLTGNWLITALLGGGFWGLFFYPGNWPIFGPTHLPLVVEGVLLSVADYTGFLYVRTGTPEYVRLIEQGSLRTFGGHTTVIAAFFSAFVSMLMYCVWWYFGKIYCTAFFYVKGERGRISMKNDVTAFGEKGFAQGIR*

>Yojoa_01_2022_E_hypo_scaffold_27567_c2_3

MSRTDEILAAAKMPPEAVRMSRYIDAVYFPILCILLVGTFHMHFMLLAGDWDFWLDWKDRQWWPVVTPIVGVMYCAALQYYLWVNYRLSYGATLCIVCLLVGEWLTRYWGFYWWSHYPLNFVLPSTMIPGALMLDTILLLTGNWLITALLGGGFWGLFFYPGNWPIFGPTHLPLVVEGVLLSVADYTGFLYVRTGTPEYVRLIEQGSLRTFGGHTTVIAAFFSAFVSMLMYCVWWYFGKIYCTAFFYVKGERGRISMKNDVTAFGEKGFAQGIR*

>Yojoa_01_2022_E_epi_scaffold_40062_c1_3

MSRTDEILAAAKMPPEAVRMSRYIDAVYFPILCILLVGTFHMHFMLLAGDWDFWLDWKDRQWWPVVTPIVGVMYCAALQYYLWVNYRLSYGATLCIVCLLVGEWLTRYWGFYWWSHYPLNFVLPSTMIPGALMLDTILLLTGNWLITALLGGGFWGLFFYPGNWPIFGPTHLPLVVEGVLLSVADYTGFLYVRTGTPEYVRLIEQGSLRTFGGHTTVIAAFFSAFVSMLMYCVWWYFGKIYCTAFFYVKGERGRISMKNDVTAFGEKGFAQGIR*

These are all the same sequence from different assemblies- when blasted to NCBI, they all hit to a Nitrosomonas and unknown Proteobacteria. There are no Nitrosomonas in Yojoa MAGs. All these genes are on scaffolds <5,000 bp. All the scaffolds only contain an A, B, C subunit of AMO with no other genes. Based on these pieces of data, we cannot place the amoA into a genome bin.

1. Castelle CJ, Hug LA, Wrighton KC, Thomas BC, Williams KH, Wu D, et al. Extraordinary phylogenetic diversity and metabolic versatility in aquifer sediment. *Nat Commun*. 2013; 4(1):2120.
2. Daims H, Lebedeva EV, Pjevac P, Han P, Herbold C, Albertsen M, et al. Complete nitrification by Nitrospira bacteria. Nature. 2015; 528(7583):504–9.
3. Rochman FF, Kwon M, Khadka R, Tamas I, Lopez-Jauregui AA, Sheremet A, et al. Novel copper-containing membrane monooxygenases (CuMMOs) encoded by alkane-utilizing Betaproteobacteria. *ISME J.* 2020; (3):714–26.
4. Tavormina PL, Orphan VJ, Kalyuzhnaya MG, Jetten MSM, Klotz MG. A novel family of functional operons encoding methane/ammonia monooxygenase-related proteins in gammaproteobacterial methanotrophs. *Environmental Microbiology Reports*. 2011; 3(1):91–100.
