## Supplementary Text 2 for "Dominant nitrogen metabolisms of a warm, seasonally anoxic freshwater ecosystem revealed using genome resolved metatranscriptomics"

**RNA preparation summary**

Briefly, rRNA was removed from 10 ng of total RNA using Qiagen FastSelect probe sets for bacterial, yeast, and plant rRNA depletion (Qiagen) with RNA blocking oligo technology. The fragmented and rRNA-depleted RNA was reverse transcribed to create first strand cDNA using Illumina TruSeq Stranded mRNA Library prep kit (Illumina) followed by second strand cDNA synthesis which incorporates dUTP to quench the second strand during amplification. The double stranded cDNA fragments were then A-tailed and ligated to JGI dual indexed Y-adapters, followed with an enrichment of the library by 13 cycles of PCR. The prepared libraries were quantified using KAPA Biosystems' next-generation sequencing library qPCR kit and run on a Roche LightCycler 480 real-time PCR instrument. Sequencing of the flowcell was performed on the Illumina NovaSeq sequencer following a 2x150nt indexed run recipe.

Full JGI protocols available at <https://jgi.doe.gov/user-programs/pmo-overview/protocols-sample-preparation-information/>
